# A novel cryopreservation and biobanking strategy to study lymphoid tissue stromal cells in human disease

**DOI:** 10.1101/2023.02.06.525604

**Authors:** Joshua D Brandstadter, Angelina De Martin, Mechthild Lütge, Antonio Ferreira, Brian T Gaudette, Yves Stanossek, Shumei Wang, Michael V Gonzalez, Edward Camiolo, Gerald Wertheim, Bridget Austin, David Allman, Megan S Lim, David C Fajgenbaum, Jon C Aster, Burkhard Ludewig, Ivan Maillard

## Abstract

Non-hematopoietic lymph node stromal cells (LNSCs) regulate lymphocyte trafficking, survival, and function for key roles in host defense, autoimmunity, alloimmunity, and lymphoproliferative disorders. However, study of LNSCs in human diseases is complicated by a dependence on viable lymphoid tissues, which are most often excised prior to establishment of a specific diagnosis. Here, we demonstrate that cryopreservation can be used to bank lymphoid tissue for the study of LNSCs in human disease. Using human tonsils, lymphoid tissue fragments were cryopreserved for subsequent enzymatic digestion and recovery of viable non-hematopoietic cells. Flow cytometry and single-cell transcriptomics identified comparable proportions of LNSC cell types in fresh and cryopreserved tissue. Moreover, cryopreservation had little effect on transcriptional profiles, which showed significant overlap between tonsils and lymph nodes. The presence and spatial distribution of transcriptionally defined cell types was confirmed by in situ analyses. Our broadly applicable approach promises to greatly enable research into the roles of LNSC in human disease.

## Introduction

Despite comprising less than five percent of cells in lymphoid tissues, non-hematopoietic lymph node stromal cells (LNSCs) have an out-sized role in shaping immune cell responses and homeostasis (1). LNSCs include heterogenous fibroblast and endothelial cell populations with profound effects on lymphocyte trafficking, survival, and activation (2) with roles in host defense, autoimmunity, alloimmunity, the tumor microenvironment, responsiveness to immune checkpoint inhibition, and inflammatory disorders (3–13).

Lymph nodes (LN) and other secondary lymphoid organs function as specialized sentinel sites for the processing of foreign antigens and the generation of specific adaptive immune responses (1, 2, 8, 14). Proper LN function requires a delicate choreography between antigenpresenting cells, T cells, B cells, and other cellular partners. LNSCs coordinate the interaction of these multiple cells via compartmentalization of regional LN microenvironments that control lymphocyte recruitment, survival, and function. LNSCs include distinct cell types, including fibroblastic stromal cells (FSCs), lymphatic endothelial cells (LECs), and blood endothelial cells (BECs). FSCs are themselves a collection of heterogenous cells with distinct immunological functions whose inflammation-induced remodeling is critical to the adaptive immune response (1, 2, 8, 14). FSC subsets include contractile and lymphocyte-interacting groups. Contractile fibroblasts include vascular smooth muscle cells (VSMCs). Immune-interacting fibroblasts include T cell zone reticular cells (TRCs) that can be further subdivided by their microanatomical location and level of expression of the chemokine CCL19, as well as perivascular reticular cells (PRCs) and B cell-interacting reticular cells (BRCs), including follicular dendritic cells (FDCs) in germinal centers. Recent single-cell transcriptomic data have heightened our understanding of lymphoid stromal cell subsets (13–19), and capturing this rich information is essential to understand the role and regulation of these cells in human disease.

Unlike hematopoietic cells that have been analyzed extensively with the aid of large-scale biobanking, the study of LNSCs in human disease has been limited. Embedded in collagen and other extracellular matrix proteins, LNSCs require enzymatic digestion to be extracted efficiently from tissues. These limitations have restricted the study of these cells in human disease to fresh tissue processed as soon as possible after harvest, a labor-intensive strategy that is costly and subject to batch effects, and by necessity often involves work-up of tissues prior to establishment of a definitive diagnosis (11, 13, 18–21). Thus, an effective cryopreservation method is needed for large-scale biobanking of lymphoid tissues to learn more about the role of LNSCs in human disease.

Here, we introduce the use of whole-tissue cryopreservation for the study of LNSCs by flow cytometry and single-cell RNA sequencing (scRNAseq). Following this protocol, we demonstrate that 2-3 mm fragments of lymphoid tissue can be cryopreserved with good subsequent recovery of viable non-hematopoietic cells, enabling the identification of the same variety of LNSC subsets by flow cytometry and scRNAseq as observed in fresh tissue. We demonstrate effective tissue cryopreservation using two different DMSO-containing reagents and following different protocols, with superior viability and cell yield compared to when cryopreservation is performed after enzymatic digestion. Our strategy can facilitate the collection of a wide variety of clinical samples without excessive upfront financial and labor commitments. Our approach also provides time for investigators to define the pathological diagnosis ahead of committing resources to a sample and to process multiple samples concurrently, thus minimizing batch effects. Altogether, this strategy enables systematic studies of LNSC cells in lymphomas and other human lymphoid tissue disorders via large-scale biobanking of lymphoid tissue.

## Results

### Tissue cryopreservation preserves cell viability to allow biobanking and subsequent isolation of lymphoid tissue stromal cells

Tissue cryopreservation would represent a major advance to enable large-scale collection of lymphoid tissues for the study of LNSCs in human disease. We developed a workflow to process cryopreserved lymphoid tissue into a single-cell suspension for flow cytometry-based immunophenotyping, cell sorting, and scRNAseq (**Figure 1A**). To systematically validate our approach, we applied our workflow to hyperplastic human tonsils received fresh as excess discarded tissue. Tonsils were cut into 2-3 mm fragments that were stirred to ensure random sampling from all anatomical parts of the tonsil (**Figure 1B**). Fragments were cryopreserved through various strategies described below and thawed or processed fresh (Figure 1A-B). These cells were then analyzed by flow cytometry or flow sorted to enrich for CD45^-^EpCAM^-^ stromal cells before scRNAseq.

**Figure 1.**
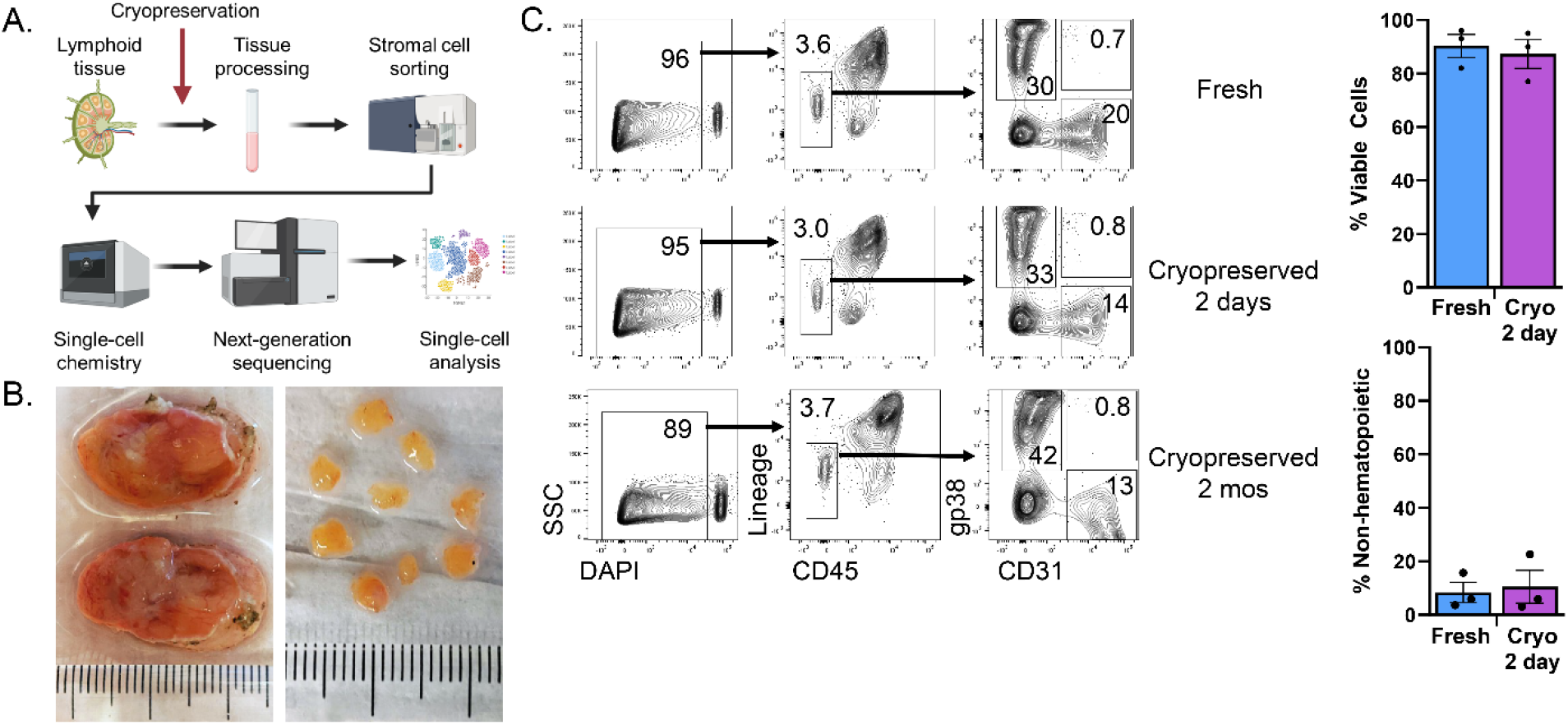
Whole tissue cryopreservation preserves cell viability to allow biobanking of lymphoid tissue. **(A)** Schematic overview of the procedure to process lymphoid tissue for single-cell RNA sequencing. Tissue processing involved gross dissection of adipose and connective tissue, cutting samples into 2-3 mm tissue fragments, and serial enzymatic digestion with collagenase, dispase, and DNase to extract LNSCs into single-cell suspensions. Selection of LNSCs by cell sorting involved gating on live singlets with a CD45^-^EpCAM^-^ immunophenotype. 10,000 sorted cells were then loaded into the 10X Chromium Controller before next-generation sequencing. **(B)** Fresh tonsils received as pathology excess specimen (top) and after gross dissection and cutting into 2-3 mm pieces (bottom). As pictured, 8-10 pieces (ca. 150mg total) were placed in each cryovial containing 1mL DMSO-containing cryopreservative for freezing **(C)** Tonsil pieces were either kept fresh in FBS-containing RPMI at 4°C for two days or cryopreserved for two days or cryopreserved for two months. Flow cytometry analysis of freshly processed, 2-day cryopreserved, or 2-month cryopreserved tonsil cells. The first column shows flow cytometry plots with live/dead staining based upon DAPI uptake. Live cells were then analyzed for expression of CD45 and hematopoietic lineage markers (CD3, CD14, CD16, CD19, CD20, CD56) with gating on non-hematopoietic cells. These non-hematopoietic cells were analyzed for expression of the fibroblast marker podoplanin (gp38) and endothelial marker (CD31), with subsequent gating identifying fibroblastic stromal cells (gp38^+^CD31^-^), blood endothelial cells (gp38^-^CD31^+^), and lymphatic endothelial cells (gp38^+^CD31^+^). Bar graphs compare viability and recovery of non-hematopoietic cells from tonsils that were freshly processed or cryopreserved for 2 days (p = 0.68 and p = 0.78, respectively, by student’s t-test, n= 3 patients/group).

We first assessed the viability of cells recovered by enzymatic digestion from cryopreserved versus fresh tonsils and key phenotypic features of the CD45^-^ stromal cell compartment (**Figure 1C**). A fraction of the processed tonsillar fragments was frozen in a DMSO-containing cryopreservative reagent (Cryostor CS10), while another fraction was stored at 4°C in RPMI medium with 2% FBS. Two days later, cryopreserved samples were thawed and digested in parallel to fresh samples kept at 4°C. The single-cell suspension was then stained and analyzed by flow cytometry. Additional cryopreserved tissue was stored at −80°C for two months before being thawed, digested, stained, and analyzed. There was a comparable proportion of viable cells in samples that had been left fresh or cryopreserved for 2 days (n = 3 patients, p = 0.68) or 2 months (**Figure 1C**). Among viable cells, fresh and cryopreserved tissue contained a similar proportion of CD45^-^ non-hematopoietic cells (p = 0.78). Furthermore, similar proportions of gp38^+^CD31^-^ FSCs among all CD45^-^ cells were recovered from fresh vs. cryopreserved tissue (30% fresh vs. 33% cryopreserved). The same was true for gp38^-^CD31^+^ BECs (20% fresh vs. 14% cryopreserved) and gp38^+^CD31^+^ LECs (0.7% fresh vs. 0.8% cryopreserved). Altogether, these data indicate that viable fibroblasts and endothelial cells could be efficiently recovered after cryopreservation of whole lymphoid tissue.

### Single-cell RNA sequencing of sort-purified CD45^-^EpCAM^-^ cells identifies distinct stromal cell subsets in fresh lymphoid tissue

To capture the diversity of stromal elements in human tonsils, we turned to scRNAseq as a sensitive and unbiased approach to identify LNSC subsets. We first generated scRNAseq data for freshly acquired tonsillar LNSCs. We processed hyperplastic tonsils into 2-3 mm fragments. After enzymatic digestion and cell sorting of CD45^-^EpCAM^-^ cells, we analyzed 27,608 total cells from three patients with tonsillar hyperplasia. The relative expression of established marker genes for FSCs (*PDGFRA, PDGFRB, CXCL13, APOE, CCL21, CCL19*, and *PDPN*), BECs (*CDH5, ENG, CD34, PECAM1*), and LECs (*PROX1, PECAM1, PDPN*) was displayed on a heat map to identify stromal cell types (**Figure 2A**). FSCs could be further sub-divided into distinct subsets (**Figure 2B-D**), including VSMCs, *ACTA2^+^* PRCs, and *CCL19^+^* TRCs. VSMCs are contractile fibroblasts that express high levels of *ACTA2* in addition to *MCAM* and genes encoding proteins involved in contractility (*TAGLN, MYH11, TPM2). ACTA2^+^* PRCs express a lower level of *ACTA2* and other contractile genes in addition to *CCL19* and collagen mRNA produced by TRCs (*COL1A1, COL1A2*). Immune-interacting TRCs could also be identified via expression of *CCL19* and *CCL21*. A subset of *CXCL13-* expresssing FDCs that sustain germinal center responses was also observed (21). These fibroblast and endothelial subset identifications are comparable to similar recently published data after accounting for divergent sorting strategies employed upstream of scRNAseq (11, 13, 18, 19). A recent detailed analysis of human tonsils identified PDPN^+^ FRCs that are comparable to our CCL19-expressing TRCs, CD21^+^ FDCs, and an *ACTA2^+^* “double-negative” (PDPN^-^CD21^-^) fibroblast subset similar to our *ACTA2^+^* PRCs and VSMCs (11).

**Figure 2.**
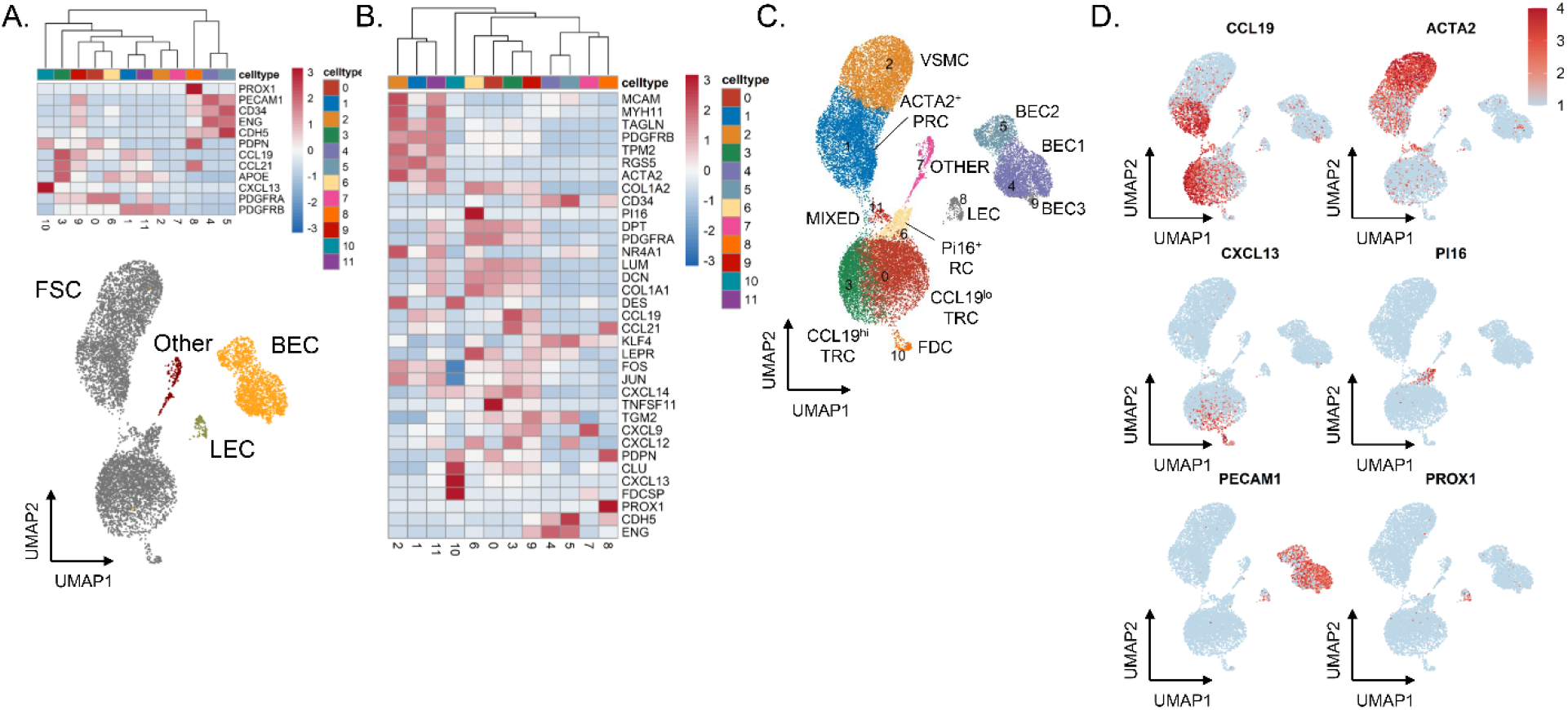
Single-cell RNA sequencing of sorted CD45^-^EpCAM^-^ cells identifies distinct stromal cell subsets in fresh lymphoid tissue. **(A)** Heatmap showing gene expression of known markers for fibroblastic stromal cells (*PDGFRA, PDGFRB, CXCL13, APOE, CCL21, CCL19*, and *PDPN*), blood endothelial cells (*CDH5, ENG, CD34, PECAM1*), and lymphatic endothelial cells markers (*PROX1, PECAM1, PDPN*). Underneath, UMAP shows all cells through Seurat-based clustering of sorted CD45^-^EpCAM^-^ cells acquired from three freshly processed tonsils after filters for quality control and removal of residual hematopoietic and epithelial cells. Cell coloring based on cell type: fibroblastic stromal cells in grey, blood endothelial cells in yellow, and lymphatic endothelial cells in green, and other cells in red. **(B)** Heatmap showing gene expression of known markers for fibroblastic stromal cell subsets, including *ACTA2^+^* perivascular reticular cells (*ACTA2*, *TAGLN, TPM2, PDGFRB*), vascular smooth muscle cells (*ACTA2*, *MYH11, MCAM*), CCL19^hi^ T-zone fibroblastic reticular cells (*CCL19*, *CCL21*, *CXCL12*, *CXCL9*), CCL19^lo^ T-zone fibroblastic reticular cells (*LUM, DCN, PDPN, PDGFRA*), Pi16^+^ reticular cells (*PI16, LEPR*), and follicular dendritic cells (*CXCL13, CLU, FDCSP, DES*). **(C)** UMAP showing Seurat-based clustering with labeled cell-types based on expression of known markers. **(D)** Feature plots show relative expression of cluster-defining markers.

### Whole tissue cryopreservation preserves diverse stromal cell subsets identified by single-cell RNA sequencing

Given the sensitivity of scRNAseq to identify LNSC subsets, we tested whether all subsets identified in fresh samples could be recovered in cryopreserved tissue. Hyperplastic tonsils were cut into 2-3 mm pieces and either stored at 4°C or cryopreserved for two days before processing. Tissue from the same tonsil was also left cryopreserved for two months before thawing and similar processing. The number of genes in each cell (nFeature_RNA), the number of molecules in each cell (nCount_RNA), and the mitochondrial content for each cell (percent.mt) were similar for tonsillar tissue processed fresh (median values 956, 1841, 5.2%, respectively), after cryopreservation for two days (median values 876, 1683, 5.9%, respectively), or after cryopreservation for two months (median values 1086, 2142, 6.0%, respectively) (**Supplementary Figure 1A**). Gene expression in tissue processed fresh or cryopreserved (averaging expression from 2-day and 2-month cryopreserved samples) was largely within a standard deviation, indicating a lack of any significant effect of cryopreservation on transcript representation (**Figure 3A**). Similar proportions of fibroblasts (grey; 77% fresh vs. 83% cryopreserved), BECs (yellow; 20% fresh vs. 14% cryopreserved), and LECs (green; 0.7% fresh vs. 0.4% cryopreserved) were recovered across fresh and cryopreserved tissue (**Figure 3B**). Furthermore, no fibroblast or endothelial cell subset was lost with cryopreservation and similar proportions of all subsets were maintained (all spatialFDR > 0.99) (**Figure 3C**, **Supplementary Figure 2A**).

**Figure 3.**
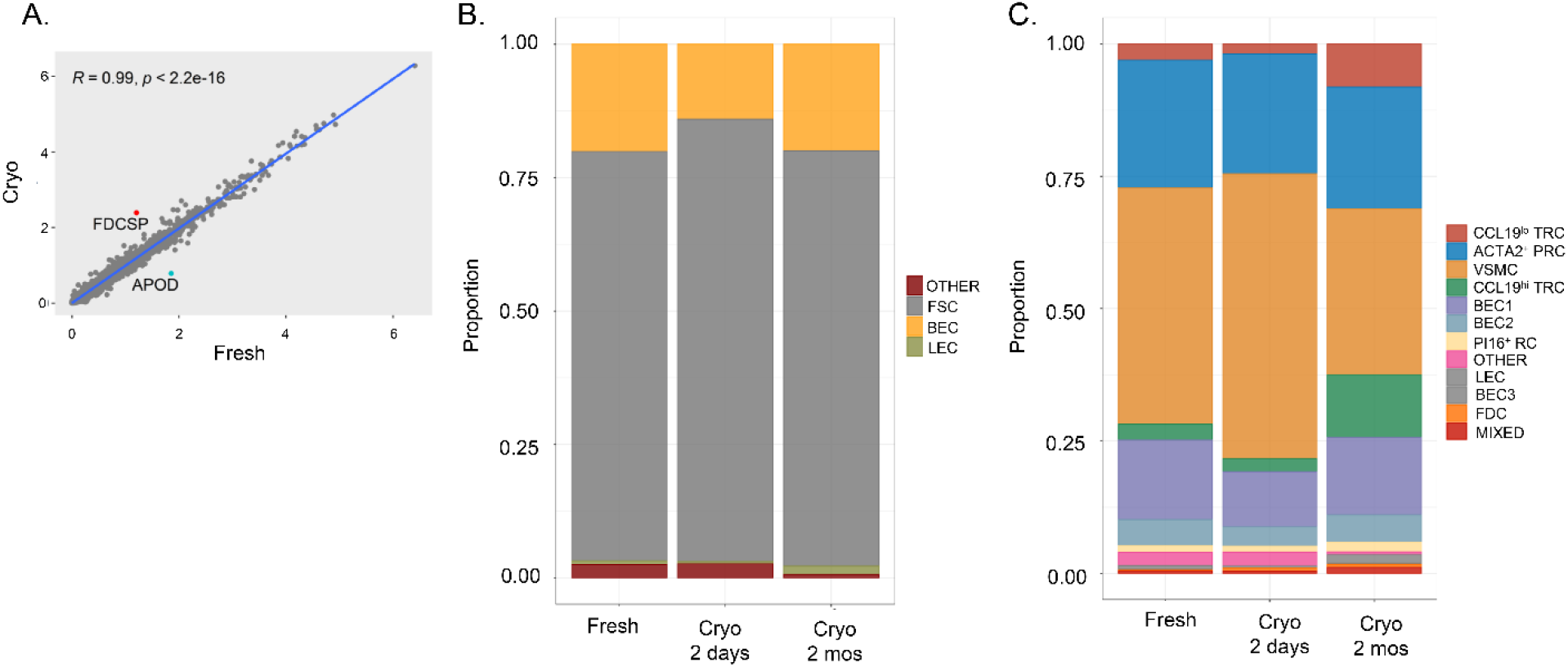
Whole tissue cryopreservation preserves diverse stromal cell subsets detected by single-cell RNA sequencing. Tonsil pieces were either kept fresh in FBS-containing RPMI at 4°C for two days vs. cryopreserved for two days or cryopreserved for two months. **(A)** Linear regression of gene expression between freshly processed and cryopreserved tonsil (with “cryo” defined as an average expression of two-day and two-month cryopreserved tissue). Pearson correlation with associated p-value listed in graph with genes up-regulated in cryopreserved tissue highlighted in red and down-regulated genes highlighted in blue. **(B)** Colors in each bar define proportion of each subset within the entire sample with fibroblastic stromal cells (FSCs) highlighted in grey, blood endothelial cells (BECs) highlighted in yellow, lymphatic endothelial cells (LECs) highlighted in green, and otherwise unidentified cells (other) highlighted in red. **(C)** Colors in each bar define proportion of each Seurat-defined cluster within the entire sample. Cluster identities are determined by expression of known markers from analysis of freshly processed tonsil in Figure 2.

### Whole tissue cryopreservation is feasible with different DMSO-containing reagents

Whole tissue cryopreservation appears successful at recovering all fibroblast and endothelial cell subsets from lymphoid tissue using a commercial DMSO-containing cryopreservative reagent, Cryostor CS10 (Sigma Aldrich, St. Louis, MO). We then asked whether we could achieve similar results using 10% DMSO and 90% FBS for cryopreservation. Cryopreservation with Cryostor reagent or DMSO/FBS were similarly successful at maintaining cell viability (77% Cryostor vs. 80% DMSO) and recovery of FSCs (gp38^+^CD31^-^; 53%vs. 56%), BECs (gp38^-^CD31^+^; 13% vs. 13%), and LECs (gp38^+^CD31^+^; 1.8% vs. 1.4%) as assessed by flow cytometry (**Supplementary Figure 3A**). We then analyzed CD45^-^EpCAM^-^ cells sorted from these conditions by scRNAseq. Samples processed fresh or through either cryopreservative strategy showed comparable quality control parameters (**Supplementary Figure 1B**). No transcript appeared to be enriched beyond a standard deviation in cryopreserved tissue compared to fresh tissue after averaging gene expression from both cryopreservation strategies (**Supplementary Figure 3B**). Expression of only one gene was enriched in Cryostor-preserved cells compared to DMSO/FBS-preserved cells – the hemoglobin beta globin gene (*HBB*), representing a likely contaminant. We observed similar recovery of a dominant FSC population (grey; 69% Cryostor vs. 75% DMSO) in addition to BECs (yellow; 23% Cryostor vs. 20% DMSO) and LECs (green) (**Supplementary Figure 3C**; 6.8% Cryostor vs. 3.2% DMSO) between cryopreserved samples using either Cryostor or DMSO in FBS. All Seurat-defined clusters were present at similar proportions in freshly processed tonsil, Cryostor-preserved tonsil, or DMSO/FBS-preserved tonsil (all spatialFDR > 0.99) (**Supplementary Figure 2A, Supplementary Figure 3D**). Altogether, our data show that several DMSO-containing cryopreservation reagents (including an inexpensive non-commercial preparation) can be used to biobank whole lymphoid tissue for the study of stromal cells.

### Cryopreservation of enzymatically digested cells impairs viability but preserves stromal cell subsets

We next studied the impact of enzymatic digestion prior to cryopreservation. The percentage of viable cells was decreased when enzymatically digested cells were cryopreserved (65%) compared to fresh tissue (93%) or whole tissue cryopreservation (90%) (**Figure 4A**). After accounting for this loss of viability, similar proportions of FSCs (59% whole tissue vs. 59% enzymatically digested cells), BECs (5% whole tissue vs. 13% enzymatically digested cells), and LECs (1.2% whole tissue vs. 0.7% enzymatically digested cells) were observed by flow cytometry. Using scRNAseq, no change in the number of gene transcripts per cell, number of molecules per cell, or increase in the mitochondrial content was observed when enzymatic digestion preceded cryopreservation (**Supplemental Figure 1C**). Similarly, no change in transcript representation by over one standard deviation was observed between freshly processed tissue or cryopreserved tissue (with averaged data from the two cryopreservation strategies) (**Figure 4B**). Whole tissue cryopreservation and cryopreservation of enzymatically digested cells also generated similar transcriptomic data within a standard deviation. FSCs (74% whole tissue vs. 86% enzymatically digested), BECs (22% whole tissue vs. 12% enzymatically digested), and LECs (1.1% whole tissue vs. 0.9% enzymatically digested) were extracted in similar proportions regardless of whether whole tissue or enzymatically digested cells were cryopreserved (**Figure 4C**). All fibroblast and endothelial cell subsets were preserved in similar proportions with cryopreservation of enzymatically digested cells compared to fresh tissue or whole tissue cryopreservation (all spatialFDR > 0.99) (**Figure 4D**, **Supplementary Figure 2A**). Thus, for samples that are large enough to tolerate loss of cell viability, creation of cell suspensions prior to cryopreservation appears to be a suitable alternative approach, although it is more labor-intensive upfront.

**Figure 4.**
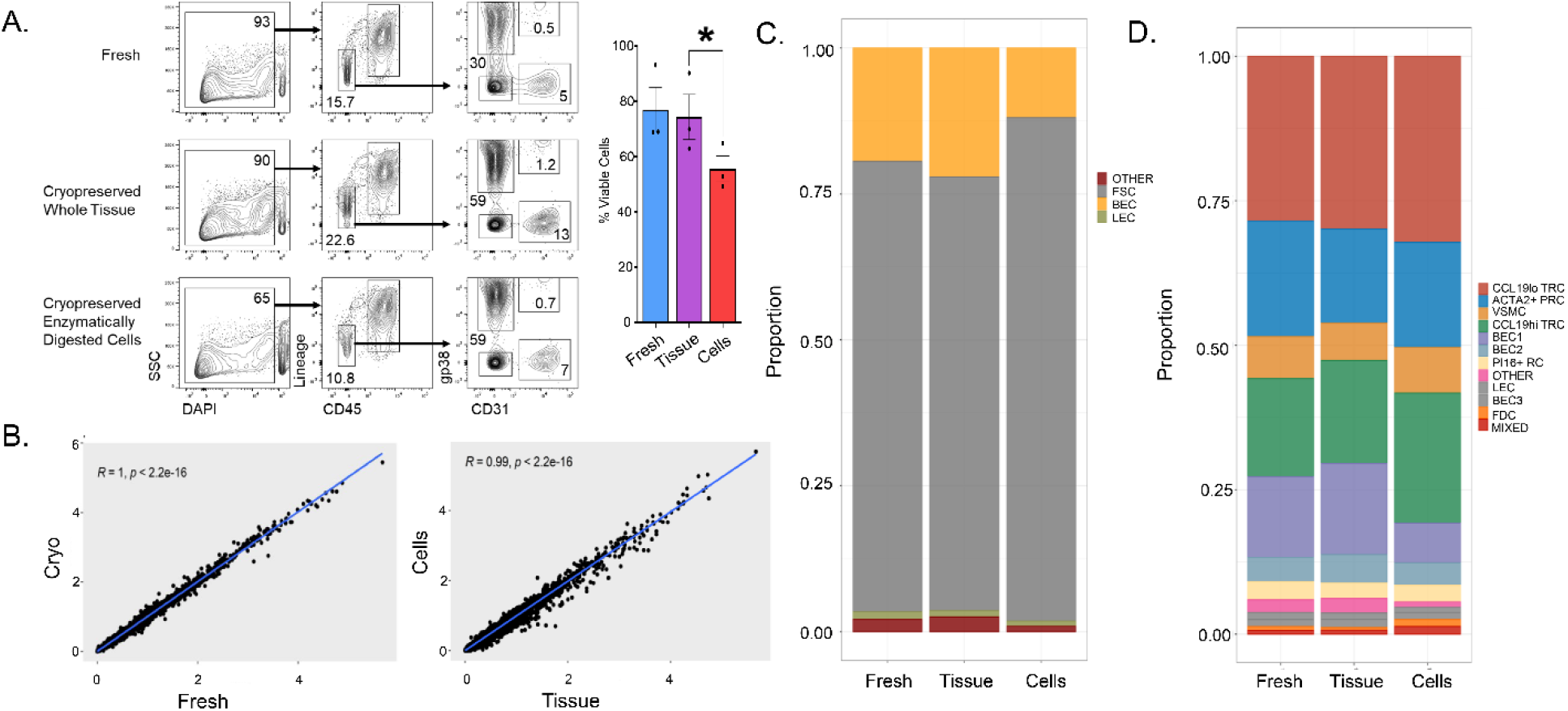
Cryopreservation of enzymatically digested cells impairs viability but preserves stromal cell subsets. Tonsil fragments were either kept fresh in FBS-containing RPMI at 4°C for two days or cryopreserved either as whole tissue or cells that have been enzymatically digested. **(A)** Flow cytometry analysis of freshly processed tonsil cells, whole-tissue cryopreserved cells, or cryopreserved enzymatically digested cells. The first column represents live/dead staining by DAPI uptake on all singlets. Viability is also shown as a bar graph (n = 3 tonsils) * p < 0.05 by student’s T test comparison after one-way ANOVA. Live cells were then analyzed for expression of CD45 and hematopoetic lineage markers (CD3, CD14, CD16, CD19, CD20, CD56) in the second column with gating showing non-hematopoietic cells. These non-hematopoietic cells were then analyzed for expression of the fibroblast marker podoplanin (gp38) and endothelial marker (CD31) with gating showing fibroblastic stromal cells (gp38^+^CD31^-^), blood endothelial cells (gp38^-^CD31^+^), and lymphatic endothelial cells (gp38^+^CD31^+^). **(B)** Linear regression of gene expression between (left) freshly processed and cryopreserved tonsil (with “cryo” defined as an average expression of whole tissue and enzymatically digested cell cryopreservation). Linear regression of gene expression between (right) whole tissue and enzymatically digested cell cryopreservation is also shown. Pearson correlation with associated p-value listed in graph. **(C)** Colors in each bar define the proportion of each subset within the entire sample with fibroblastic stromal cells (FSCs) highlighted in grey, blood endothelial cells (BECs) highlighted in yellow, lymphatic endothelial cells (LECs) highlighted in green, and otherwise un-identified cells (other) highlighted in red. **(D)** Colors in each bar define proportion of each Seurat-defined cluster within the entire sample. Cluster identities are determined via expression of known markers from analysis of freshly processed tonsil in Figure 2.

### In situ localization of lymphoid stromal cell subsets identified by single-cell RNA sequencing

To assess if gene expression data from cryopreserved lymphoid tissue could be used for downstream spatial analysis, we stained tissue sections for specific proteins and transcripts. Our scRNAseq analysis of sorted CD45^-^EpCAM^-^ cells from fresh tonsils identified major populations of contractile and immune-interacting fibroblasts (**Figure 2**). Contractile fibroblasts express VSMC genes, suggesting a role as perivascular cells. Immune-interacting fibroblasts express chemokines, such as CCL19 and CCL21, and other molecules consistent with their identity as fibroblastic reticular cells (FRCs) that form fine conduits for soluble antigen transport and other immunological functions. *ACTA2^+^* PRCs were predicted to act as FRCs based on prior work and their collagen and chemokine expression profile(2). We sought to verify these histologic inferences about the fibroblast subsets identified in our scRNAseq data by immunofluorescent microscopy of tonsillar tissue. Cells that stain for alpha-smooth muscle actin (aSMA, the *ACTA2* gene product) were observed to encircle CD31^+^CD34^+^ endothelial cells in tonsils (**Supplementary Figure 4A-C**) consistent with the expected distribution of VSMCs. In contrast, cells expressing CCL19 or another FRC marker, podoplanin (PDPN), showed a reticular morphology and were scattered diffusely throughout the tissue. Another group of ACTA2-expressing cells were observed to form mesh-like reticular networks, fitting with their identity as ACTA2^+^ PRCs (**Supplementary Figure 4B**). To further validate the microanatomical positioning of our transcriptionally defined subsets, we employed in situ hybridization to directly detect mRNA. As proof-of-principle, our scRNAseq data identified an FDC subset of fibroblasts that expressed *FDCSP* mRNA. Consistent with our transcriptomic data, we detected *FDCSP* mRNA in a germinal center pattern, while *CCL19* mRNA was present in interfollicular regions (**Supplementary Figure 4D**).

Altogether, this work demonstrates that whole tissue cryopreservation can be employed to facilitate the study of LNSCs in human disease. Despite concern that cryopreservation would impair cell yield or gene expression due to the need for enzymatic digestion to extract LNSC, we could recover cells with comparable viability. Furthermore, whole tissue cryopreservation – including with a simple 10% DMSO solution in fetal bovine serum – led to no loss of even rare LNSC subsets and no global impact on transcript representation. Features of LNSCs identified in tonsils by flow cytometry and single-cell RNA sequencing could be orthogonally confirmed by fluorescent microscopy and in situ hybridization.

### Tonsillar stromal cells have LN correlates and model LN tissue

Given limited access to fresh lymphoid tissue, we used hyperplastic tonsils to test the efficacy of cryopreservation strategies. However, LNs are the lymphoid tissue of primary clinical interest – the most common microenvironment for lymphomas and other lymphoproliferative disorders. We therefore sought to compare tonsils to LNs to determine commonalities and differences in their stromal composition. Following the same experimental protocol, we processed three LNs that had non-specific inflammatory (“reactive”) histologies as compared to freshly processed tonsils. We generated scRNAseq data from a total of 20,397 analyzable LN cells. Analysis of all freshly processed tonsillar and LN cells in a single dimensionality reduction UMAP showed significant overlap between clusters regardless of tissue of origin with the exception of one cluster that was predominantly observed in tonsils and a nearby cluster disproportionately composed of LN cells (**Figure 5A**). Using relative expression of established markers for FSCs, BECs, and LECs (**Supplementary Figure 5A**), we determined the cell type of Seurat-defined clusters (**Figure 5B**). Overall, we observed similar proportions of each FSCs (77.1%±0.9% tonsil vs. 75.6%±10.6% LN), BECs (18.9%±1.5% tonsil vs. 19.5%±10.6% LN), and LECs (1.8%±1.3% tonsil vs. 4.3%±0.7% LN) across samples irrespective of tissue of origin. Similarly, we used the relative expression of fibroblast subset markers (**Supplementary Figure 5B**) to identify Seurat-defined clusters (**Figure 5C**, **Supplementary Figure 5C**). Comparing the proportions of each LNSC subset, we observed distinct tonsillar (LogFC = 6.00, SpatialFDR = 0.11, p = 0.051) and LN (LogFC = −7.83, SpatialFDR = 0.089, p = 0.032) CCL19^+^ TRC clusters and increased Pi16+ PRCs in LN tissue compared to tonsil (LogFC = −5.67, SpatialFDR = 0.094, p = 0.038) (**Figure 5C**, **Supplementary Figure 2B**). While tonsil and LN subsets expressed *CCL19* similarly, they differentially expressed other genes, such as *CCL21, CXCL12, TNFSF11*, and *CXCL9* (**Supplementary Figure 5B-E**). Altogether, these data suggest that LN CCL19^+^ TRC have a more inflammatory phenotype compared to the corresponding tonsillar cells, possibly as a result of sampling LN with reactive inflammation. Otherwise, similar cell-type representation and gene expression was observed upon comparing tonsillar to LN stromal cells.

**Figure 5.**
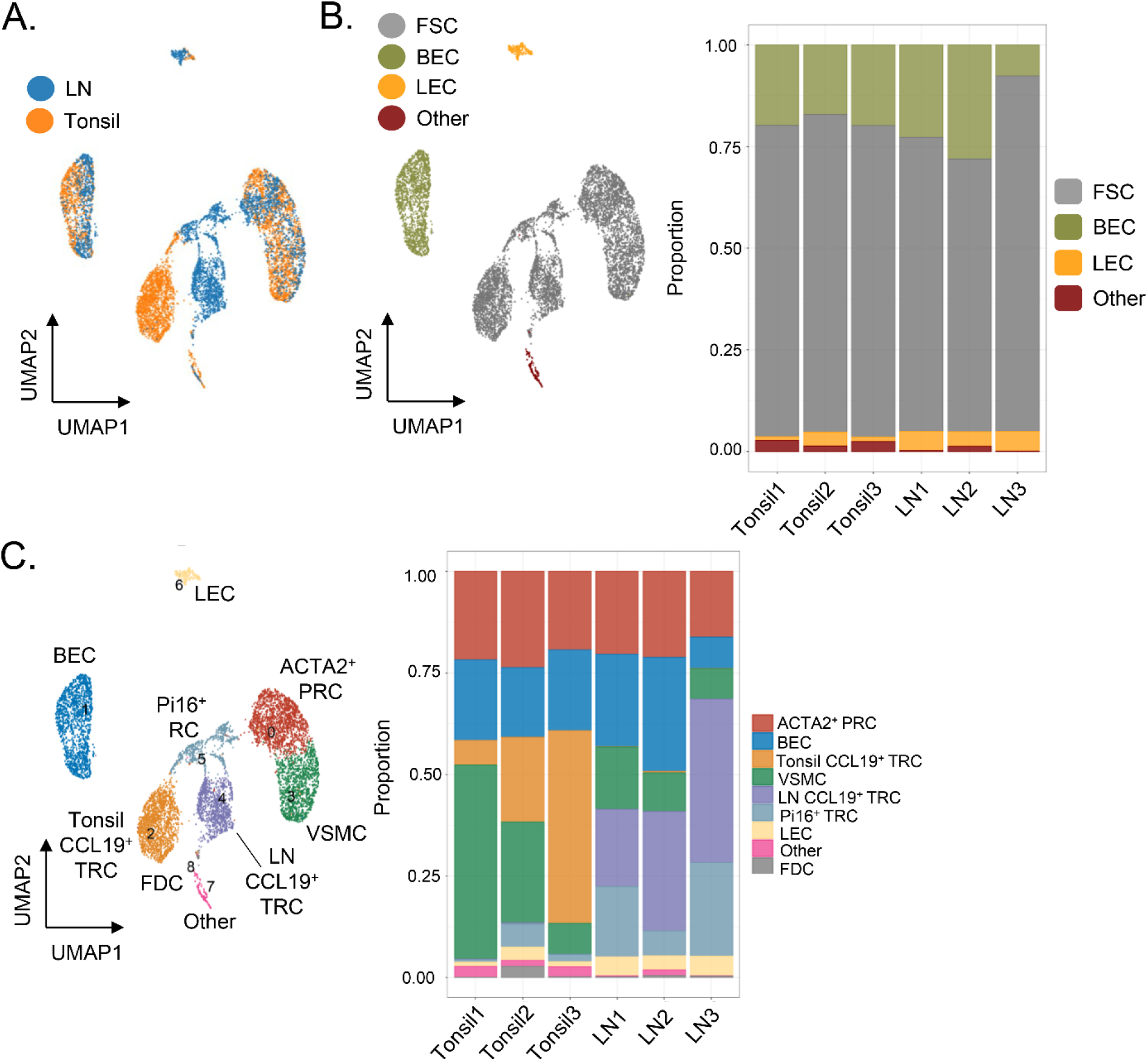
Tonsillar stromal cells have LN correlates and model LN tissue. Three freshly processed hyperplastic tonsils were compared to three LNs with reactive histologies following the schematic in Figure 1A. **(A)** UMAP shows all cells through Seurat-based clustering of sorted CD45^-^EpCAM^-^ cells after filters for quality control and removal of residual hematopoietic and epithelial cells with coloring based on lymphoid tissue type. **(B)** Cell-types were identified by relative gene expression observed in Supplementary Figure 2A. UMAP (left) is colored based on cell-type identities. Relative proportion of each cell-type shown for each sample is shown in bar graph (right) with colors defining proportions fibroblastic stromal cells (FSCs) in grey, blood endothelial cells (BECs) in yellow, lymphatic endothelial cells (LECs) in green, and otherwise un-identified cells (other) in red. **(C)** Cell-types in UMAP (left) are labeled according to relative gene expression observed in Supplemental Figure 2B. Bar graph (right) with colors in each bar defining proportion of each Seurat cluster.

## Discussion

We have established a versatile approach for whole lymphoid tissue cryopreservation that can allow biobanking of specimens and build insights into how lymphoid tissue stromal cells influence human health. By testing alternative cryopreservation strategies, we showed that small pieces of lymphoid tissue frozen in a broadly available, non-commercial solution can enable long-term biobanking of clinical samples despite the added stress of enzymatic digestion needed to recover LNSCs. Altogether, our insights pave the way to a scale of research that will be needed to establish the roles of LNSCs in human disease.

So far, a heavy dependence on fresh tissue limited the investigation of LNSCs in human disease. For example, follicular lymphoma – a common hematological malignancy where stroma has long been suggested to influence lymphomagenesis and chemotherapy refractoriness – was recently shown to involve remodeling of LNSCs (11). However, this study was limited to 3-4 follicular lymphoma LNs that could be processed as fresh tissue, without comparison to other types of lymphomas or to reactive LNs. Instead, mechanistic insights into human lymphoid stromal cells were achieved using more readily available tonsils, a limitation in the applicability of the findings noted by the authors. Other recent work described a novel LNSC niche that plays a key role in dendritic cell homeostasis in mice and that was shown to be critical in T cell immunity (19). Confirmation of the applicability of this finding to humans relied upon data from only 3 resting human LNs resected during cancer staging surgeries, where a LNSC subset with a similar transcriptomic signature was observed. The availability of a biobanking approach as described here will enable collection of large numbers of LNs, which will both control for the heterogeneity inherent in clinical samples and facilitate the study of rare or unusual LN pathologies.

Peripheral T cell lymphomas such as angioimmunoblastic T cell lymphoma (AITL) and atypical lymphoproliferative disorders such as Castleman disease have characteristic histologic aberrations in LNSCs that make them attractive for further investigation. For AITL, this includes an increase in FDC numbers, a highly arborized vasculature, expanded B cell subsets and heterogeneous inflammatory cells (22). Histologic aberrations in stroma and an expansion of non-neoplastic immune cells can complicate the diagnosis of AITL. Castleman disease is characterized by prominent FDCs in addition to hypervascularity and atretic germinal centers (23). In addition, past work suggested clonality of LNSCs in the unicentric form of Castleman disease (24–26). Insights into the role that LNSCs might play in their pathogenesis will be facilitated by cryopreservation strategies and could inform the development of new diagnostic approaches for these rare diseases with a challenging diagnosis.

We found superior stromal cell viability for whole tissue cryopreservation compared to cryopreservation of enzymatically digested cells, although analysis of all viable cells found no change in transcript representation or loss of any LNSC subset. Other strategies such as single nuclei RNA sequencing can be useful to extract single cell transcriptomic information from complex tissues, including tissue cryopreserved shortly after harvest. Indeed, this approach can be used in place of single-cell RNA sequencing to avoid the challenges of extracting cells from extracellular matrix (27, 28). Single nuclear RNA sequencing protects against dissociation-induced gene alterations and can detect rare cell-types missed by other RNA-sequencing strategies (28–30). Despite these advantages, all LNSCs are rare cell types in bulk lymphoid tissue and single nuclear sequencing without dissociation into single-cell suspension would prohibit LNSC enrichment by CD45 bead- or column-based depletion or cell sorting. The loss of LNSC enrichment would prevent detailed analysis of LNSCs as most of the sample would be comprised of hematopoietic cells. Particularly rare but important LNSC subtypes – such as FDCs that are critical for germinal center responses – would be especially difficult and costly to capture with single-nuclear RNA sequencing alone, as it would mandate RNA sequencing from a very large number of nuclei. Therefore, whole tissue cryopreservation remains the more attractive approach for the study of LNSCs.

Hyperplastic tonsils, typically discarded without pathologic review after resection, provided a large quantity of regularly available tissue to evaluate whether cryopreserved and fresh tissues could produce comparable quality data. However, LNs are the lymphoid tissue that harbors the vast majority of lymphomas and other lymphoproliferative disorders and where further investigation of LNSCs would be of most clinical interest. We therefore, compared tonsillar to LN stroma using LNs with non-specific inflammatory (“reactive”) histologies. We found substantial overlap of fibroblastic and endothelial cell populations despite tissue of origin. The greatest distinction between the tissues was CCL19^+^ TRC subset, which appears more inflammatory in LNs with higher levels of CCL21, CXCL9, CXCL12, and TNFSF11. Overall, this more inflammatory phenotype to an immune-interacting subset observed in LN compared to tonsillar stroma fits with the reactive inflammatory phenotype observed histologically in reactive LNs.

Altogether, we have developed whole tissue cryopreservation to advance the study of LNSCs in human disease by providing a method whereby tissue can be banked for later study. We hope that this will foster insights into how LNSCs might contribute to the lymphoma microenvironment, atypical lymphoproliferative disorders such as Castleman Disease, responsiveness to immune checkpoint inhibitor treatment as well as other infectious and autoimmune diseases. Additional technical advances, including use of spatial transcriptomic approaches that might better define LNSC-lymphocyte interactions, will provide even greater insight into the potential roles these cells play in human disease.

## Materials and Methods

### Human tonsil collection and gross dissection

De-identified, anonymous human tonsils were procured as pathology excess from pediatric donors at the Children’s Hospital of Philadelphia via a material transfer agreement with the University of Pennsylvania. Patients were six to 21 years of age with tonsillar hyperplasia. Samples were received on the same day as surgical resection. Using a scalpel blade, electrocauterized fragments and adipose tissue were carefully removed, and tonsils were then cut finely into 2-3 mm pieces. Specifically, tonsils were cut longitudinally with parallel incisions to create thick sections that were then successively cut across the perpendicular axis into smaller and smaller pieces until each piece was 2-3 mm in size. Smaller fragments maximize the surface area subsequently exposed to enzymes, thus enabling faster and more complete digestion. Tonsil fragments were mixed by gentle stirring to ensure uniform sampling from all areas of the tonsil. Finally, the pieces were stored fresh at 4°C in RPMI-1640 media with 10% fetal bovine serum (FBS) or cryopreserved as described below.

### Human LN collection

Patients with lymphadenopathy undergoing surgical resection provided consent to donate a portion of their resected LN for research per protocol approved by the University of Pennsylvania Institutional Review Board (Protocol #826185). A portion of all lymph nodes was fixed in formalin for hematopathology review of a hematoxylin and eosin-stained slide by a single experience hematopathologist (M.S.L.) to identify reactive histology.

### Whole-tissue cryopreservation and thawing

First, tonsils were cut with a scalpel into 2-3 mm pieces as described above. Next, fragments were gently stirred to prevent selection biases from overrepresentation of any part of the tonsil. After mixing, 8-10 pieces (ca. 150mg total) were placed in each cryovial containing 1mL Cryostor CS10 (Sigma Aldrich, St. Louis, MO) or 1mL 90% FBS/10% DMSO and placed in Corning CoolCell LX Cell Freezing Container (Sigma Aldrich) for storage in a −80°C freezer. Cryotubes were thawed in 37°C water baths and samples were transferred to 5mL FACS tubes where they were washed three times in RPMI with 2% FBS before addition of enzymatic digestion solution. At least three cryovials were used to generate sufficient digested cells so that 10,000 sorted CD45^-^ EpCAM^-^ cells could be loaded into the Chromium Controller (see below).

### Enzymatic digestion of tonsils

Fresh or thawed cryopreserved samples were incubated in 37°C water bath with 3mL enzymatic digestion solution. Enzymatic digestion of fresh or thawed, cryopreserved tissue was performed otherwise similarly to prior descriptions (31, 32). An enzymatic medium was prepared by adding 2 mg/mL dispase II (Gibco), 0.6 mg/ml collagenase P (Sigma Aldrich), and 0.3 mg/mL DNase I (Sigma Aldrich) to pre-warmed RPMI with 2% FBS and 20mM HEPES. Every 10 minutes, samples were agitated by pipetting using a 1000μl micropipette. Every 15 minutes, the digestion medium with cells in suspension was aspirated, transferred through 70μm filter to 20mL ice-cold FACS buffer (PBS containing 2% FBS, 20mM HEPES, and 2mM EDTA) before being spun at 1500rpm for 5 minutes and resuspended in fresh FACS buffer. Fresh digestion medium was immediately added to the residual tissue for a total of three serial, 15-minute digestions.

### Flow Cytometry Antibodies

For flow cytometry, the following antibodies were used: anti-CD45 APCFire750 (clone HI30, Biolegend,1:100), anti-EpCAM APCFire750 (clone 9C4, Biolegend, 1:100), anti-podoplanin PE-Cy7 (clone NC-08, Biolegend, 1:200), and anti-CD31 PerCP Cy5.5 (clone WM59, Biolegend, 1:200). DAPI (Sigma Aldrich) at 10μg/mL final concentration was added immediately before sorting for live-dead cell exclusion.

### Low-pressure flow cytometry and cell sorting

Flow cytometric cell sorting was performed with the FACSDIVA software on a BD FACSAria (Franklin Lakes, NJ) set to 20 psi with a 100μm nozzle. The sorting protocol was as previously described, although we did not require any bead-based enrichment prior to sorting and could compensate successfully with antibody-stained UltraComp eBeads Compensation Beads (Invitrogen, Waltham, MA) (31). We sorted ≥ 30,000 CD45^-^ EpCAM^-^ live singlet cells into 1.5mL DNA LoBind tubes (Eppendorf, Hamburg, Germany) containing 40μl FBS. Purity was routinely > 95%. Tubes were spin-concentrated with only a part of the supernatant removed by gentle aspiration to leave ~50μl volume in the tube. The concentrated cells were then fully resuspended in the remaining volume and counted with a hemacytometer using a Trypan Blue viability dye. Flow cytometric analysis was performed using FlowJo v.10.6.1 (Ashland, OR).

### Single-cell RNAseq library preparation and sequencing

Sorted cells were loaded into a 10X Chromium Controller and next-generation sequencing libraries were built using Chromium Next GEM Single Cell 3’ GEM, Library, & Gel Bead Kit v3.1 kit according to manufacturer’s instructions (10X Genomics, Pleasanton, CA). After sorting, 10,000 cells were loaded per sample into the Chromium Controller and captured in gel bead emulsions. The mRNA was reverse transcribed and the cDNA was amplified for 11 cycles with manufacturer-supplied primers. Libraries were then constructed through fragmentation, adapter ligation, and a sample indexing PCR with double-sided size selection using SPRIselect beads (Beckman Coulter, Brea, CA). All samples were quantified by Qubit High-sensitivity DNA fluorometry (Thermo Fisher Scientific, Waltham, MA) and checked for library quality by TapeStation High-sensitivity D5000 (Agilent Technologies, Santa Clara, CA). Indexed samples were pooled, denatured, and diluted to 1.8pM before being loaded onto a NextSeq 500/550 High Output Kit v2.5 (150 cycles, Illumina, San Diego, CA) for paired-end sequencing on a NextSeq 550 (Illumina).

### Bioinformatic analysis

Raw sequencing files were aligned to the Ensembl human GRCh38 reference genome in Cell Ranger software (v.3.1.0, 10X Genomics) (33). Quality control was performed in R (v.4.2.0) with the Seurat R package (v.4.1.1) in order to remove damaged cells or doublets/multiplets based on high or low UMI counts (filtering out feature counts over 3,000 or less than 200) or high percent mitochondrial genes (filtering out cells with > 15% mitochondrial counts) (34). Cells that had positive read counts between 200 and 3000 and percent mitochondrial genes less than 15% were retained. Cells expressing leukocyte-specific (*CD3E, PTPRC, CD79A, MZB1, CCR7, CD7, CD52*) or epithelial marker genes (*KRT5, KRT15*, and *KRT17*) or *MKI67* were also removed. After quality control to remove damaged and contaminating cells, the final dataset included 22,691 human non-hematopoietic tonsillar cells across samples for downstream analysis. Downstream analysis was performed with the Seurat R package (v.4.1.1) with normalization, scaling, dimensionality reduction with PCA and UMAP, and clustering (34). Clusters were labeled based on calculated cluster-defining gene expression and known markers reported in previous publications (2, 16, 35).

Following cluster identification, individual samples were compared based on how the upstream tissue was processed (fresh vs. different cryopreservation strategies as described above). This comparison was done by determining the proportion of each cluster that accounts for the total cells in each sample. In this way, differentially abundant cell-types that may have weathered cryopreservation relatively poorly or better could be identified compared with cells from freshly processed samples. Additionally, the Milo framework for differential abundance testing was employed (36, 37). The *milo* package constructs a KNN graph (k = 30, p = 0.1) to assign cells to neighborhoods distinct from Seurat-defined clusters. Individual neighborhoods were assigned cell-type labels based on Seurat-defined clusters on the basis of majority voting of the cells in that neighborhood.

### Immunofluorescence microscopy

Tissues were embedded in FSC 22 Clear (Leica Biosystems), fresh frozen in an isopropanol-dry ice bath and stored at −80°C. 10 μm sections were cut with a cryostat (Leica CM1950) and mounted onto Thermo slides and fixed for 10 min in methanol at −20°C. Mounted tissues were then blocked with PBS containing 10% FCS, 1 mg/ml anti-Fcγ receptor (BD Biosciences) and 0.1% Triton X-100 (Sigma) at 4°C for 2 hours. These slides were incubated overnight with anti-human CD34 FITC (Clone 581, Biolegend 343504, 1:200), anti-human/mouse ACTA2 Cy3 (Clone 1A4, Sigma C6198, 1:1000), anti-human CD31 A647 (Clone WM59, Biolegend 303112, 1:100) or unconjugated anti-human CCL19 (polyclonal, R&D AF361-SP, 1:200) that was detected with anti-goat IgG A488 (Jackson Immunoresearch 705-545-003, 1:1000) and unconjugated anti-human PDPN (Clone NZ-1.3, eBioscience 14-9381-82, 1:200) that was detected with anti-rat IgG A647 (Jackson Immunoresearch 712-605-153, 1:1000) secondary antibodies. Microscopy was performed using a confocal microscope (LSM-980, Carl Zeiss), and images were recorded and processed with ZEN 2010 software (Carl Zeiss). Imaris Version 9 (Bitplane) was used for image analysis.

### RNA-Scope

Formalin-fixed paraffin-embedded tissues were sectioned to 5 μm using a microtome (Leica RM2255). Assays for *CCL19* and *FDCSP* mRNA (Catalog 474361 and 444231, respectively, Advanced Cell Diagnostics) were used according to the manufacturer’s RNA-Scope TM Multiplex Fluorescent v2 kit instructions (Advanced Cell Diagnostics). Images were obtained using a confocal microscope (LSM-780, Zeiss) and images were recorded and processed with ZEN 2012 software (Zeiss). ImageJ 1.49v software (Wayne Rasband) was used for image analysis, rendering, masking, and reconstruction.

### Statistics

Statistical comparisons were made using one-way ANOVA and student’s t-tests where p < 0.05 was considered significant using GraphPad Prism. Linear regression and Pearson correlations were calculated using the *stats* and *ggplot2* packages, respectively, in the R programing environment. To compare differential gene expression, a linear regression was fitted to data comparing the normalized log2-transformed gene expression of the comparison groups. A gene was considered differentially expressed if it was one standard deviation off the fitted line as defined an absolute residual > 1. Statistics related to differential abundance analysis were made using the *milo* package in the R programming environment(36). Neighborhoods were considered differentially abundant when spatialFDR was < 0.1, logFC > or < 0, and Benjamini-Hochberg corrected p-value < 0.05.

### Study approval

The study protocol conformed to the ethical guidelines of the 1975 Declaration of Helsinki. Tonsil use was approved by a material transfer agreement between the University of Pennsylvania and Children’s’ Hospital of Philadelphia (ID: 58590/00). All tonsils were received without any protected health information or identifiers, exempt from review by the Children’s Hospital of Philadelphia Institutional Review Board (SOP 407, Section XII “Secondary Use of De-Identified Data or Specimens”). All lymph nodes were received after patient provided consent as part of a University of Pennsylvania Institutional Review Board protocol (#826185).

### Data sharing statement

All de-identified data generated or analyzed are available in the main text, supplementary materials, or were deposited to be publicaly available through the Gene Expression Omnibus (accession GSE224661, https://www.ncbi.nlm.nih.gov/geo/query/acc.cgi?acc=GSE224661).

## Abbreviations

LNSC: Lymph node stromal cell
LN: Lymph node
VSMC: Vascular smooth muscle cell
TRC: T cell zone reticular cell
PRCs: Perivascular reticular cell
BRC: B cell-interacting reticular cell
FDC: Follicular dendritic cell
scRNAseq: Single-cell RNA sequencing

## Acknowledgements

J.D.B. is supported by a Doris Duke Charitable Foundation’s Physician Scientist Fellowship and American Society of Transplantation and Cellular Therapy New Investigator Award. J.D.B. previously received support from the American Society of Clinical Oncology Young Investigator Award and American Society of Hematology Research Training Award for Fellows. Additional support was from NHLBI (T32-HL07439 to J.D.B.) and the NCI (T32-CA009140 to B.T.G.). This work was supported by grants from the National Institutes of Health (R01-AI091627 to I.M., R01-HL141408 to D.C.F.).

## Author Contributions

J.D.B. was responsible for experimental design, performing experiments, and preparing the manuscript. A.M. and Y.S. performed immunofluorescence microscopy. M.L. and M.V.G. provided bioinformatic support. A.F. and S.W. performed in situ hybridization microscopy. B.G. provided support with performing experiments and bioinformatics. E.C. and G.W. helped with tonsil acquisition and regulatory approval. M.S.L. reviewed pathology of all lymph nodes. B.L., J.C.A., D.C.F, and D.A. contributed to experimental design and manuscript preparation. I.M. was responsible for experimental design and preparing the manuscript. All authors reviewed the final manuscript.

