## Supplementary Figures 1 - 5 for "A novel cryopreservation and biobanking strategy to study lymphoid tissue stromal cells in human disease"

**Supplementary material**

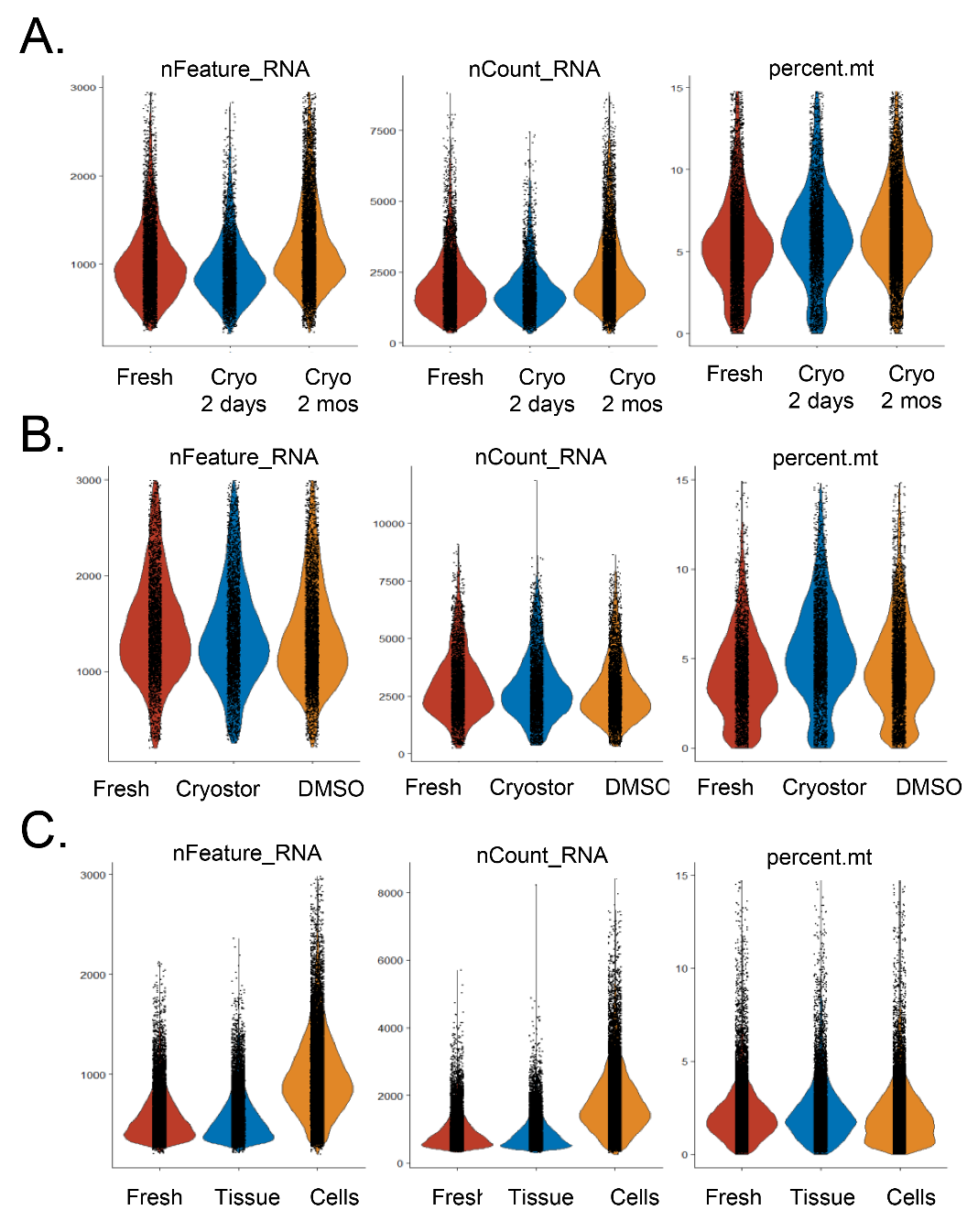
**Supplementary Figure 1. Different cryopreservation strategies show no difference in single-cell RNA sequencing quality metrics.** Violin plots show the number of genes in each cell (nFeature_RNA), the number of molecules in each cell (nCount_RNA), and the mitochondrial content for each cell (percent.mt). **(A)** Comparison of samples processed fresh, after cryopreservation for two days, or after cryopreservation for two months. **(B)** Comparison of samples processed fresh, after cryopreservation with a commercial DMSO-containing cryopreservative reagent (Cryostor), or after cryopreservation with 10% DMSO in fetal bovine serum. **(C)** Comparison of samples processed fresh, after whole-tissue cryopreservation, or after cryopreservation of enzymatically digested cells.

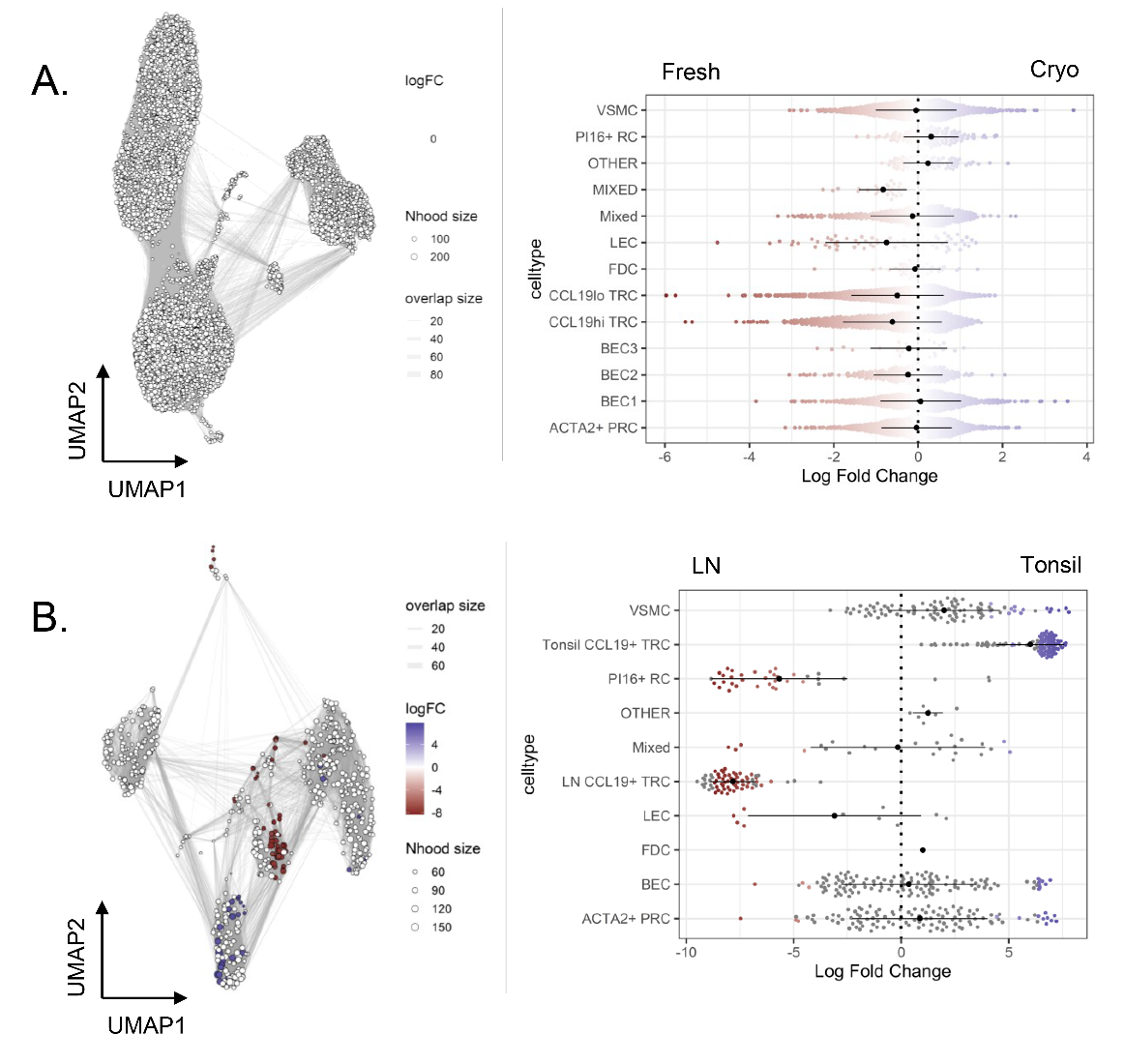
**Supplementary Figure 2. Tonsillar stromal composition is unaffected by cryopreservation but varies from LN stroma.** **(A)** *Left.* Milo neighborhoods superimposed on UMAP of CD45^-^EpCAM^-^ cells from Figure 2 where dot size correlates to number of cells in each neighborhood and line width correlates to overlap between neighborhoods. All neighborhoods are gray, indicating no differentially abundant (spatialFDR < 0.1) neighborhoods between fresh and cryopreserved tissue. *Right.* Bee-swarm plot of log-fold change (x-axis) across fresh and cryopreserved samples in Milo neighborhoods grouped together by Seurat-defined cell identity. Neighborhood needed to have ≥ 70% overlap with cell identity defined through Seurat to be included. Since all neighborhoods were not differentially abundant (spatialFDR > 0.99), all are shown on this plot (alpha = 1.0) with red indicating neighborhoods enriched in freshly processed tissue and blue indicating neihborhoods enriched in cryopreserved tissue. **(B)** *Left.* Milo neighborhoods superimposed on UMAP of tonsillar and LN cells from Figure 7 where dot size correlates to number of cells in each neighborhood and line width correlates to overlap between neighborhoods. Red indicates neighborhoods differentially abdunant (spatialFDR < 0.1) in LNs and blue for neighborhoods differentially abundant in tonsil between fresh and cryopreserved tissue. *Right.* Bee-swarm plot of log-fold change (x-axis) across fresh and cryopreserved samples in Milo neighborhoods grouped together by Seurat-defined cell identity. Neighborhood needed to have ≥ 70% overlap with cell identity defined through Seurat to be included. Only neighborhoods were differentially abundant (spatialFDR < 0.1) are shown (alpha = 1.0) with signficantly (p < 0.05) differentially abundant neighborhoods colored in red (for differentially abundant in LN) or blue (for differentially abundant in tonsil).

**Supplementary**
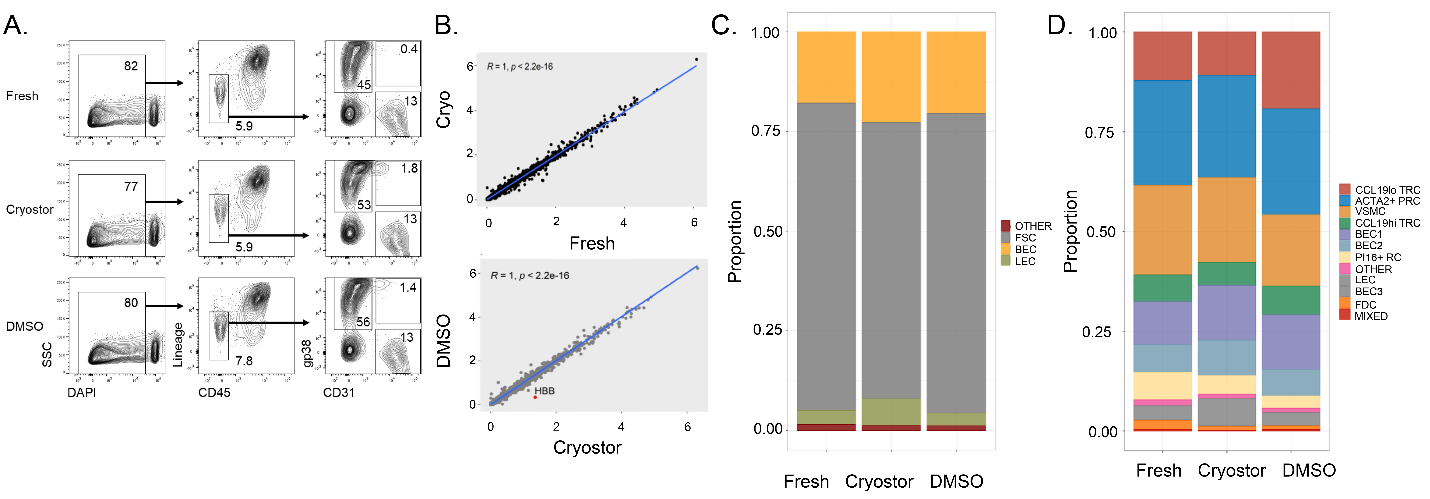
**Figure 3. Whole tissue cryopreservation is feasible with different DMSO-containing reagents.** Tonsil pieces were either kept fresh in FBS-containing RPMI at 4°C for two days or cryopreserved for two days with a commercial DMSO-containing reagent (Cryostor) or 10% DMSO in fetal bovine serum. **(A)** Flow cytometry analysis of freshly processed tonsil and tonsil tissue cryopreserved with Cryostor or 10% DMSO in fetal bovine serum. The first column represents live/dead staining by DAPI uptake on all singlets. Live cells were then analyzed for expression of CD45 and hematopoetic lineage markers (CD3, CD14, CD16, CD19, CD20, CD56) in the second column with gating showing non-hematopoietic cells. These non-hematopoietic cells were then analyzed for expression of the fibroblast marker podoplanin (gp38) and endothelial marker (CD31) with gating showing fibroblastic stromal cells (gp38^+^CD31^-^), blood endothelial cells (gp38^-^CD31^+^), and lymphatic endothelial cells (gp38^+^CD31^+^). **(B)** Linear regression of gene expression between (top) freshly processed and cryopreserved tonsil (with “cryo” referring to both Cryostor- and DMSO/FBS-preserved tissue). Linear regression of gene expression between (bottom) Cryostor- and DMSO/FBS-preserved tissue is also shown. Pearson correlation with associated p-value listed in graph with up-regulated mRNA in Cryostor-preserved tissue highlighted in red. **(C)** Colors in each bar define the proportion of each subset within the entire sample with fibroblastic stromal cells (FSCs) highlighted in grey, blood endothelial cells (BECs) highlighted in yellow, lymphatic endothelial cells (LECs) highlighted in green, and otherwise un-identified cells (other) highlighted in red. **(D)** Colors in each bar define proportion of each Seurat-defined cluster within the entire sample. Cluster identities are determined by expression of known markers as shown in Figure 2.

**Supplementary**
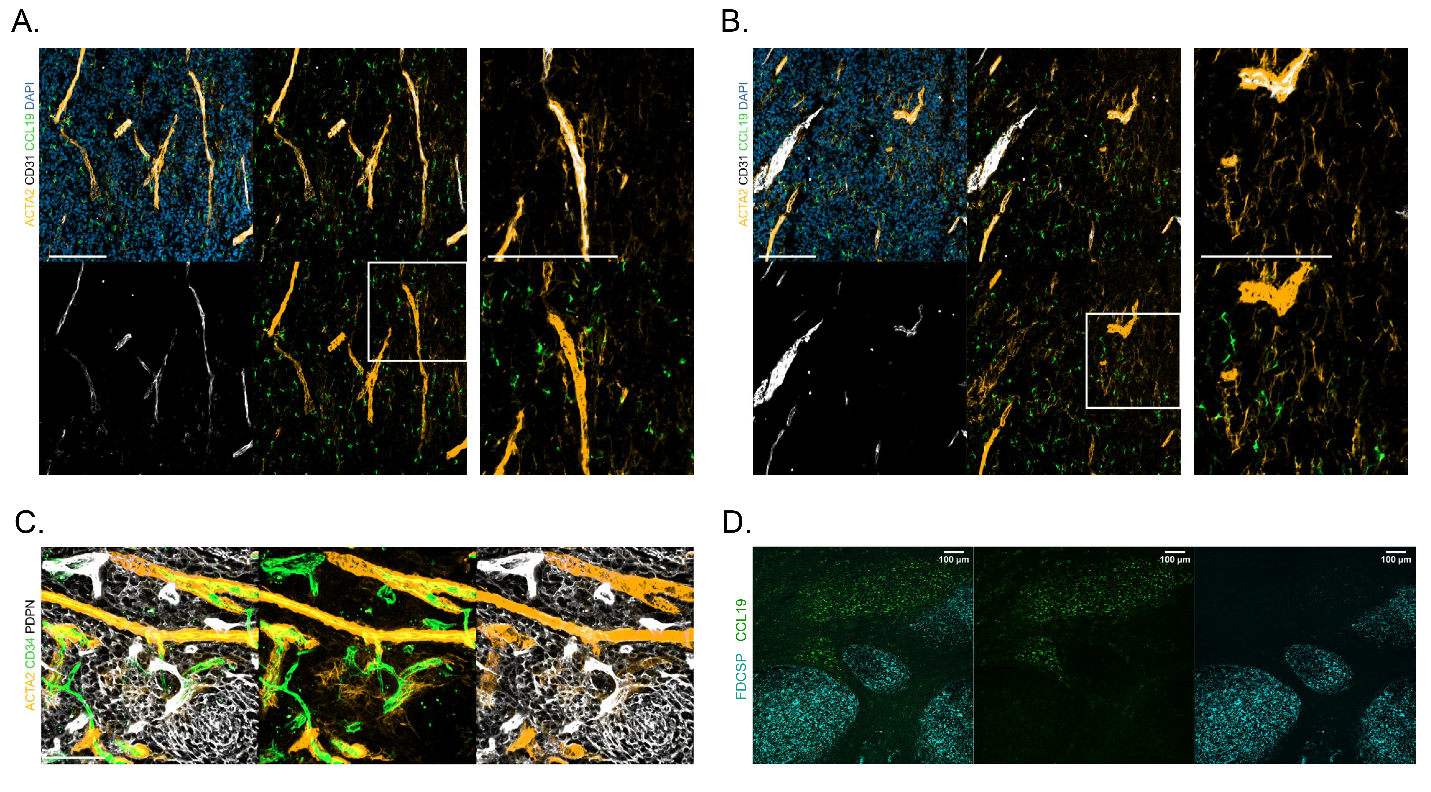
**Figure 4. Histologic appearance of lymphoid stromal cell subsets identified by single-cell RNA sequencing.** (A) Immunofluorescent microscopy with 20X objective with staining for perivascular stromal cells by aSMA/ACTA2 (yellow), endothelial cells by CD31 (white), fibroblastic reticular cells by CCL19 (green), as well as DAPI for nuclear staining (blue). **(B)** Immunofluorescent microscopy with 20X objective of the tonsil interfollicular region with the same approach. **(C)** High-resolution imaging with 40X objective showing perivascular cells by ACTA2 (yellow) encircling endothelial cells expressing CD34 (green) with separate population of fibroblastic reticular cells idetnfied via PDPN expression (white). **(D)** In situ hybridization with 25X objective of a tonsil showing germinal centers expressing the follicular dendritic cell marker FDCSP (blue) and interfollicular areas where cells that express CCL19 (green).

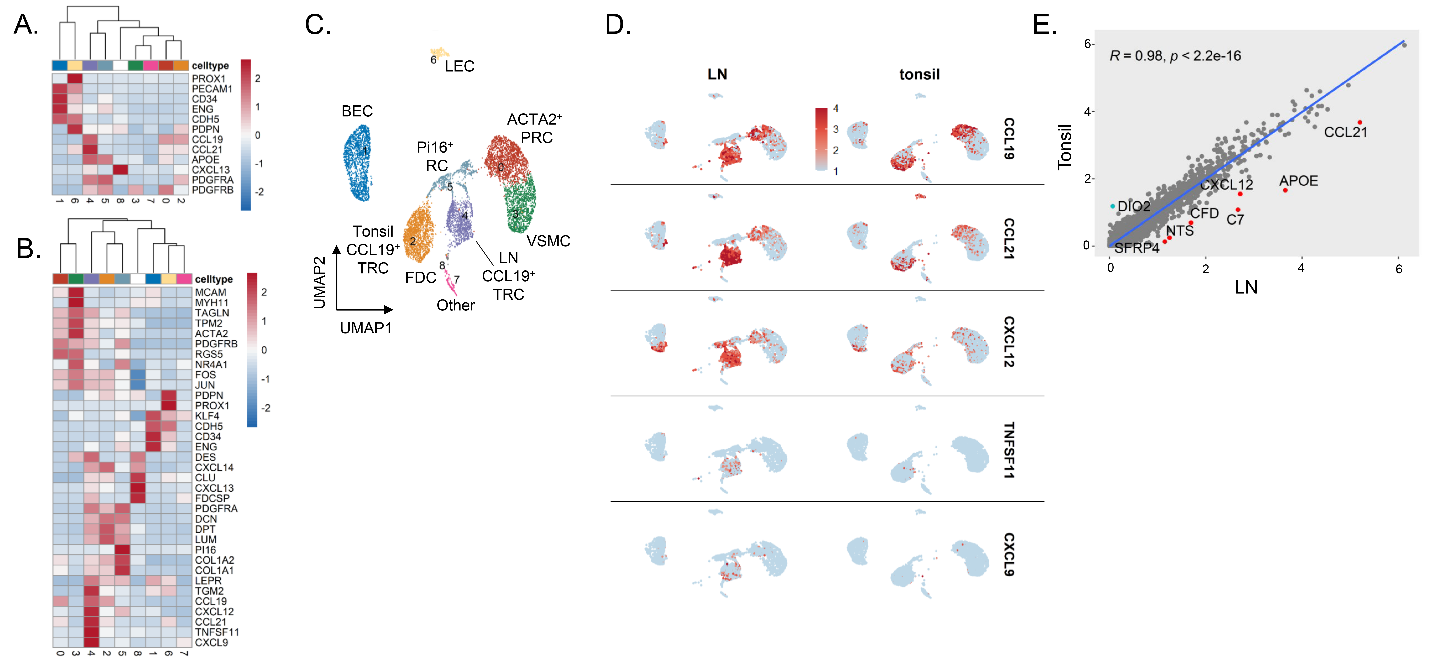
**Supplementary Figure 5. Overlapping tonsillar and LN stromal cell subsets with more inflammatory immune-interacting CCL19^+^ TRC observed in LNs.** Three freshly processed hyperplastic tonsils were compared to three LNs with reactive histologies. All cells through Seurat-based clustering of sorted CD45^-^EpCAM^-^ cells after filters for quality control and removal of residual hematopoietic and epithelial cells. **(A)** Heatmap showing gene expression for Seurat-defined clusters of known markers for fibroblastic stromal cells (*PDGFRA, PDGFRB*, *CXCL13*, *APOE*, *CCL21*, *CCL19*, and *PDPN*), blood endothelial cells (*CDH5*, *ENG*, *CD34*, *PECAM1*), and lymphatic endothelial cells markers (*PROX1*, *PECAM1*, *PDPN*). **(B)** Heatmap showing gene expression of known markers for fibroblastic stromal cell subsets, including *ACTA2*^+^ perivascular reticular cells (*ACTA2*, *TAGLN*, *TPM2*, *PDGFRB*), vascular smooth muscle cells (*ACTA2*, *MYH11*, *MCAM*), CCL19^hi^ T-zone fibroblastic reticular cells (*CCL19*, *CCL21*, *CXCL12*, *CXCL9*), CCL19^lo^ T-zone fibroblastic reticular cells (*LUM*, *DCN*, *PDPN*, *PDGFRA*), Pi16^+^ reticular cells (*PI16*, *LEPR*), and follicular dendritic cells (*CXCL13*, *CLU*, *FDCSP*, *DES*). **(C)** UMAP shows all cells through Seurat-defined clusters with labels based on subset identity derived from relative expression of known marker genes. **(D)** Feature plots show relative expression of *CCL19*, *CCL21*, *CXCL12*, *TNFSF11*, and *CXCL9* between LN-derived and tonsil-derived samples. **(E)** Linear regression of gene expression between tonsils and LNs. Pearson correlation with associated p-value listed for genes up-regulated in LNs tissue (highlighted in red) and down-regulated genes (highlighted in blue).
